# Synthesis of Cellulose Nanofiber Hydrogel Beads from *Typha angustifolia* for the Immobilization of Protease

**DOI:** 10.64898/2026.11.18.767499

**Authors:** Blessing Eleojo Ameh, Evans C. Egwim, Ifeoma F. Ossamulu

**Affiliations:** Department of Biochemistry, School of Life Sciences, Federal University of Technology Minna, Niger State, Nigeria

**Keywords:** *Typha angustifolia*, cellulose nanofiber, hydrogel beads, enzyme immobilization, protease, invasive biomass valorization

## Abstract

*Typha angustifolia*, an invasive aquatic macrophyte, poses a persistent challenge to wetland and irrigation management, yet its cellulose-rich biomass represents an underexploited feedstock for value-added biomaterials. This study reports the synthesis of cellulose nanofiber (CNF) hydrogel beads from *T. angustifolia* and evaluates their performance as a support matrix for protease immobilization. Cellulose was isolated by chlorite delignification and alkaline treatment, oxidized with sodium periodate to introduce aldehyde functional groups, blended with polyvinyl alcohol (PVA), and crosslinked with glutaraldehyde to form hydrogel beads. The beads exhibited a high water-absorption capacity of 399.67 ± 0.53% and a mechanical strength of 1.27 MPa. Covalent immobilization of crude protease onto the beads increased specific activity nearly threefold, from 24.56 ± 0.39 U/mg for the free enzyme to 59.20 ± 1.06 U/mg for the immobilized enzyme, corresponding to a loading efficiency of 50.61%. The immobilized enzyme also displayed broadened thermal and pH optima, shifting from 40 °C to 50 °C and from pH 8.0 to 9.0, respectively, and an extended operational half-life from 4.40 h to 11.37 h. These findings support the valorization of invasive *Typha* biomass into a robust, biodegradable support system with potential application in industrial biocatalysis.

## 1. Introduction

*Typha angustifolia*, commonly known as narrowleaf cattail, is a widespread aquatic plant found in wetland habitats worldwide [1]. Although the species contributes to natural filtration and sediment stabilization, its rapid vegetative growth has rendered it a problematic invasive species in numerous agricultural and ecological settings [2, 3]. Uncontrolled proliferation of *T. angustifolia* frequently obstructs irrigation channels, invades rice paddies, and displaces native biodiversity, creating substantial challenges for water management and farming [4].

Management of *T. angustifolia* is costly. Rather than treating this biomass solely as an environmental burden requiring disposal, recent advances in green chemistry favour its conversion into value-added bioproducts. This strategy serves two complementary purposes: reducing the ecological footprint of the invasion, and generating industrially useful materials [5].

The primary structural component of *T. angustifolia* is cellulose, the most abundant renewable biopolymer on Earth [6, 7]. Through chemical and mechanical processing, cellulose can be deconstructed into cellulose nanofibers (CNF) [8]. CNFs are characterized by a high aspect ratio, extensive surface area, and exceptional mechanical properties, including high strength and stiffness [8–11]. Unlike bulk cellulose, CNFs present a dense network of surface hydroxyl groups that facilitates chemical modification and interaction with other molecules [12].

However, direct utilization of raw *Typha* fibres is often limited by the presence of lignin and hemicellulose. To address this, chemical pretreatments such as chlorite delignification followed by periodate oxidation are needed to expose the cellulose fibres and introduce reactive functional groups—principally aldehydes—that enhance the material’s reactivity and binding capacity for subsequent applications [13, 14].

Among the various configurations of nanocellulose materials, CNF-based hydrogels have attracted considerable attention as support materials [15]. Hydrogels are three-dimensional, crosslinked polymeric networks that can absorb and retain large amounts of water while maintaining structural integrity [16–18]. Fabricating hydrogel beads from *Typha*-derived CNF, often reinforced with polymers such as polyvinyl alcohol (PVA), creates a matrix with tunable porosity and superior mechanical stability [14, 19]. These porous structures are particularly useful for biotechnological applications because they resemble the natural, water-based environment of biological molecules, allowing substances to move freely while still providing structural support [20, 21]. Crosslinking agents such as glutaraldehyde further stabilize these beads, preventing disintegration under operational conditions [22, 23].

One important application of these nanostructured supports is enzyme immobilization. Enzymes are powerful biological catalysts used across many industrial processes owing to their high specificity and environmentally benign reaction conditions [24, 25]. Despite their effectiveness, free enzymes have several limitations, including instability under harsh thermal and pH conditions, susceptibility to autolysis, and difficulty of recovery and reuse [26, 27]. Enzyme immobilization— the attachment of enzymes to a solid support—addresses these limitations by increasing structural stability and simplifying separation of the enzyme from the reaction mixture [27, 28]. Covalent immobilization onto nanoscale supports such as CNF is particularly effective: the high surface-area-to-volume ratio of nanofibres maximizes enzyme loading, while covalent bonding prevents enzyme leaching, improving both stability and reusability [29–31].

Proteases, a class of enzymes that catalyze the hydrolysis of peptide bonds, are among the most commercially important enzymes, holding dominant market shares in the detergent, food-processing, leather, and pharmaceutical industries [32, 33]. Industrial application of proteases, however, requires them to withstand elevated temperatures and varying pH, conditions that typically denature free enzymes [34, 35]. Developing robust immobilization strategies for proteases is therefore a priority for industrial biotechnology [36].

A range of cellulosic feedstocks—including bacterial cellulose, cotton cellulose, kenaf nanocellulose, sugarcane bagasse, rice husk, and banana pseudostem—have been explored as sources of CNF for enzyme immobilization [37, 38, 42], and each carries practical trade-offs that constrain large-scale adoption. Bacterial cellulose is highly pure but costly to produce; cotton cellulose is agriculturally valuable and therefore competes with textile and food-related demand; and lignocellulosic residues such as bagasse, rice husk, and banana pseudostem generally require more intensive delignification because of their comparatively higher lignin content. *Typha* biomass, by contrast, is neither a food nor a fibre crop and is available largely as a by-product of wetland-management efforts, so its conversion into CNF carries an environmental co-benefit— invasive-species control—that conventional cellulose feedstocks do not offer, in addition to its low or negative feedstock cost. To date, however, the specific use of CNF derived from invasive *T. angustifolia* for protease immobilization has not been widely studied. Existing work has established the potential of *Typha* biomass for bioethanol production [5], but the performance of *Typha*-derived CNF hydrogel beads as a catalytic support has not been fully elucidated, and there remains limited understanding of how the morphological properties of *Typha* CNF influence the kinetic behaviour and stability of immobilized proteases [39, 40].

We hypothesized that cellulose nanofibers isolated from *T. angustifolia* could be chemically oxidized and combined with PVA to produce hydrogel beads with sufficient porosity and mechanical integrity to support covalent protease immobilization, and that this immobilization would improve enzyme loading, catalytic performance, and operational stability relative to the free enzyme. Accordingly, this study aimed to (i) synthesize CNF hydrogel beads from *T. angustifolia*, (ii) characterize their physicochemical and structural properties, and (iii) evaluate their performance as a support for protease immobilization relative to the free enzyme.

## 2. Materials and Methods

All chemicals and reagents used were of analytical grade.

### 2.1. Isolation and Oxidation of Cellulose Nanofibers (CNF) from *Typha angustifolia*

Typha angustifolia was collected from Katsina State, Nigeria [author to confirm the exact collection site and date, as the accompanying thesis records the collection site as Maiduguri, Borno State — please reconcile before submission], and washed thoroughly under running water to remove impurities before being processed into powder. The powder was further washed several times with hot water (100 °C) to remove residual impurities.

The washed pulp was combined with sodium chlorite and acetic acid (0.6 g of sodium chlorite per gram of fibre) at a 3% solids content and subjected to delignification at 70 °C for 2 h; this step was repeated until the supernatant solution ran clear, and the bleached pulp was then rinsed thoroughly. The chlorite-delignified pulp was mixed with 9% NaOH and left to react for 1 h at 24 °C to remove residual hemicellulose [13].

For oxidation, 40 g of CNF aqueous suspension (5 wt% solids) was combined with 60 mL of distilled water and dispersed by magnetic stirring. A 50% NaIO_4_ solution was then added, and the mixture was stirred in darkness for 48 h to introduce aldehyde groups via periodate oxidation. The oxidized product was recovered by centrifugation at 23,200 × g for 12 min and rinsed with distilled water [14].

### 2.2. Synthesis of CNF Hydrogel Beads

A total of 0.60 g of oxidized CNF and 2 g of polyvinyl alcohol (PVA *[molecular weight and degree of hydrolysis not specified in the source records — author to insert manufacturer specification]*) were mixed with 30 mL of water and heated with stirring at 95 °C for 1 h to fully dissolve the PVA and disperse the CNF. After cooling to 60 °C, five drops of 0.5 M HNO3 were added as a catalyst [14]. The resulting solution was added dropwise into a 2.3 M NaOH solution to form spherical beads by ionic gelation. The beads were then immersed in 25 mL of glutaraldehyde (3% v/v) for 12 h to permit covalent crosslinking, rinsed, and dried at 60 °C [23].

The proposed reaction sequence underlying bead formation and subsequent enzyme immobilization is summarized schematically in Figure 1: hydroxyl groups on the native CNF are converted to aldehyde groups by periodate oxidation; the oxidized CNF is combined with PVA and gelled into beads; glutaraldehyde crosslinks the network and introduces additional pendant aldehyde groups; and these aldehyde groups subsequently condense with primary amine groups on the enzyme surface (principally lysine residues) to form Schiff-base (imine) linkages that covalently anchor the protease to the bead matrix.

**Figure 1.**
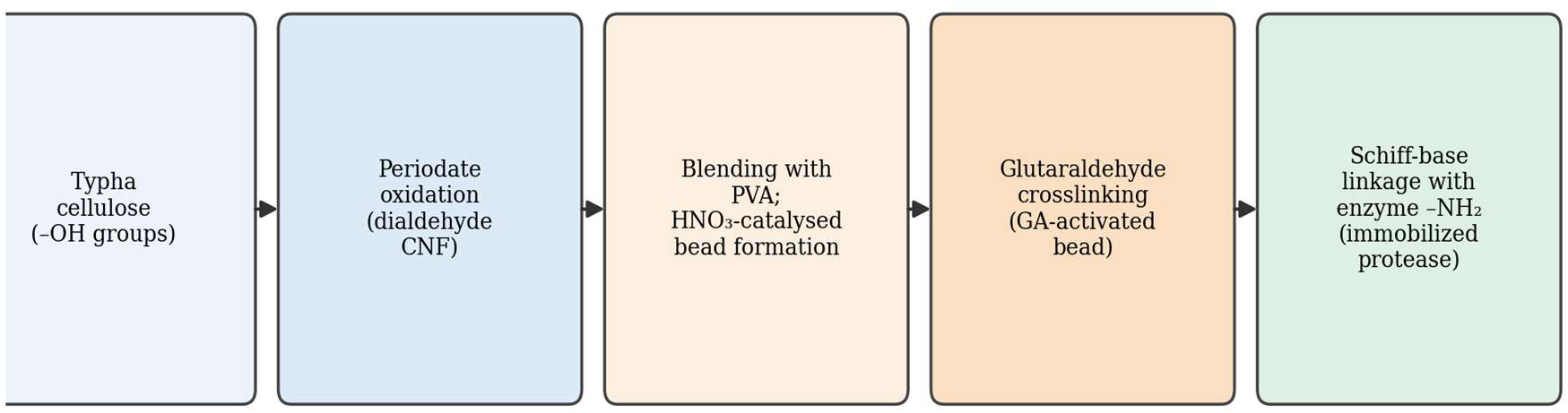
Proposed reaction scheme for the conversion of *Typha* cellulose into CNF hydrogel beads and the subsequent covalent immobilization of protease via glutaraldehyde-mediated Schiff-base formation.

### 2.3. Characterization of Beads

Swelling degree was determined by immersing 1 g of crosslinked beads in distilled water for 24 h and calculated using Equation 1 [41]:

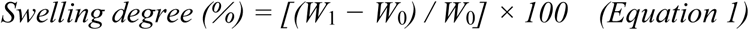

where W_0_ (g) is the dry mass and W_1_ (g) is the wet mass after 24 h.

Mechanical properties were tested by compressing beads to 60% deformation at a crosshead speed of 5 mm/min. Surface morphology was examined by scanning electron microscopy (SEM) after coating samples with a thin (5 nm) layer of gold [42].

### 2.4. Immobilization of Protease

A total of 0.10 g of glutaraldehyde-crosslinked beads and 0.5 mL of phosphate buffer (pH 6.5) were combined with 1 mL of crude protease and incubated at 25 °C for 120 min. The beads were then washed, and the protein content remaining in the supernatant was measured *[assay method — e.g., Bradford or Lowry — not specified in the source records; author to insert]* to enable calculation of loading efficiency using Equation 2:

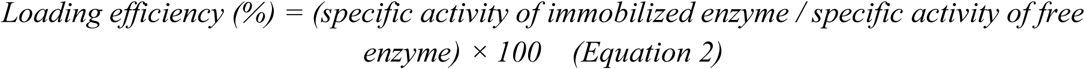

### 2.5. Protease Source

The crude protease used in this study was obtained from a dumpsite soil sample collected in Minna, Niger State, Nigeria, and was used without further purification. This crude, mixed-source preparation should be borne in mind when comparing the reported specific activities with studies employing purified, single-source proteases.

### 2.6. Enzymatic Assays and Stability

Protease activity was measured using casein as substrate (0.04–0.2%) at 60 °C and pH 6.7 for 20 min [43]. Kinetic parameters (K_m_ and V_max_) were determined from Lineweaver–Burk double-reciprocal plots. The effects of temperature (30–70 °C) and pH (5.0–10.0) on activity were evaluated for both free and immobilized enzyme preparations.

Operational stability was assessed by monitoring residual activity over a continuous 5 h assay period, from which the deactivation constant (K_d_) and operational half-life (t_1/2_) were derived using Equation 3:

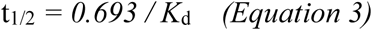

Storage stability was additionally assessed by maintaining free and immobilized enzyme preparations at 4 °C *[the storage-stability dataset supplied to us duplicates the 5-h operational-stability time course rather than presenting an independent multi-day (e.g., 7-, 30-, 60-, or 90-day) assay; the authors should substitute the genuine long-term storage dataset here, or otherwise revise Section 3.8 and Table 5 accordingly — see accompanying review response letter, Comment 9]*.

### 2.7. Statistical Analysis

All experiments were performed in triplicate, and results are expressed as mean ± standard error of the mean (SEM). Differences between free and immobilized enzyme preparations were evaluated by *[one-way ANOVA followed by Tukey’s HSD post-hoc test / Student’s t-test — author to confirm the specific test applied]* using *[statistical software and version, e.g., SPSS v.X or GraphPad Prism vX — author to insert]*, with differences considered statistically significant at p < 0.05. Superscript letters in Tables 1–5 denote statistically significant differences between means within a column.

## 3. Results and Discussion

### 3.1. Physical and Chemical Properties of CNF Hydrogel Beads

The synthesized CNF-hydrogel beads (Figure 2) exhibited a fairly uniform spherical shape with a diameter of approximately 5.0 mm when wet. Upon drying, the beads showed slight deformation from water evaporation, resulting in a rougher surface texture while retaining structural integrity [46, 47].

**Figure 2.**
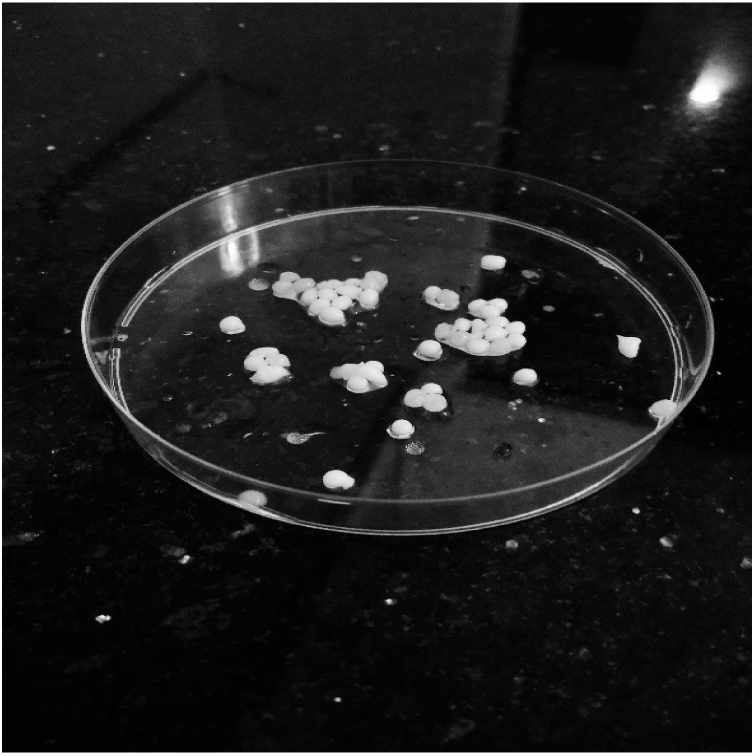
*Typha angustifolia*-derived CNF hydrogel beads.

The swelling behaviour and mechanical strength of the beads are summarized in Table 1. The beads demonstrated a swelling percentage of 399.67 ± 0.53%, indicating a substantially higher water-absorption capacity than previously reported cellulose-based beads [48]. This high water-retention capacity is advantageous for applications requiring facile substrate diffusion; conversely, very high swelling can also dilute the local density of immobilized enzyme within the bead and, in principle, compromise dimensional stability under flow, so this trade-off should be considered when translating the beads to continuous-flow bioprocessing. The measured mechanical strength (1.27 MPa) indicates a structure sufficiently robust for bioprocessing applications, comparable to, though slightly lower than, some previously reported cellulose hydrogel systems [19] and higher than others [49].

**Table 1.** Physical properties of CNF-hydrogel beads. Values are mean ± SEM of three replicates.

| **Sample** | **Swelling test (%)** | **Mechanical strength (MPa)** |
|---|---|---|
| CNF-hydrogel beads | 399.67 ± 0.53 | 1.27 |

### 3.2. Structural Characterization (FTIR)

The FTIR spectrum of the synthesized beads (Figure 3) confirms successful isolation of cellulose nanofibers from *Typha angustifolia* and their subsequent crosslinking into a stable hydrogel network [19, 53, 54]. The broad absorption band at 3301–3623 cm^-1^ corresponds to O–H stretching vibrations, confirming the hydrophilic character of the material and the extensive hydrogen bonding required for water retention. Preservation of the cellulosic backbone following delignification is indicated by characteristic peaks at 2937.1 cm^-1^ (C–H stretching) and 1140.6 cm^-1^ (C–O–C pyranose ring vibration). Successful functionalization of the nanofibres is shown by the peak at 1651.2 cm^-1^, attributed to carbonyl (C=O) groups introduced during periodate oxidation, which serve as reactive crosslinking sites. Formation of the hydrogel network is further supported by the absorption band at 1088 cm^-1^, characteristic of C–O stretching in the PVA matrix, consistent with the formation of acetal bridges between the oxidized CNF and PVA chains. Collectively, these results indicate that *Typha* biomass was successfully converted into an oxidized, crosslinked cellulose–PVA hydrogel matrix suitable for enzyme immobilization.

**Figure 3.**
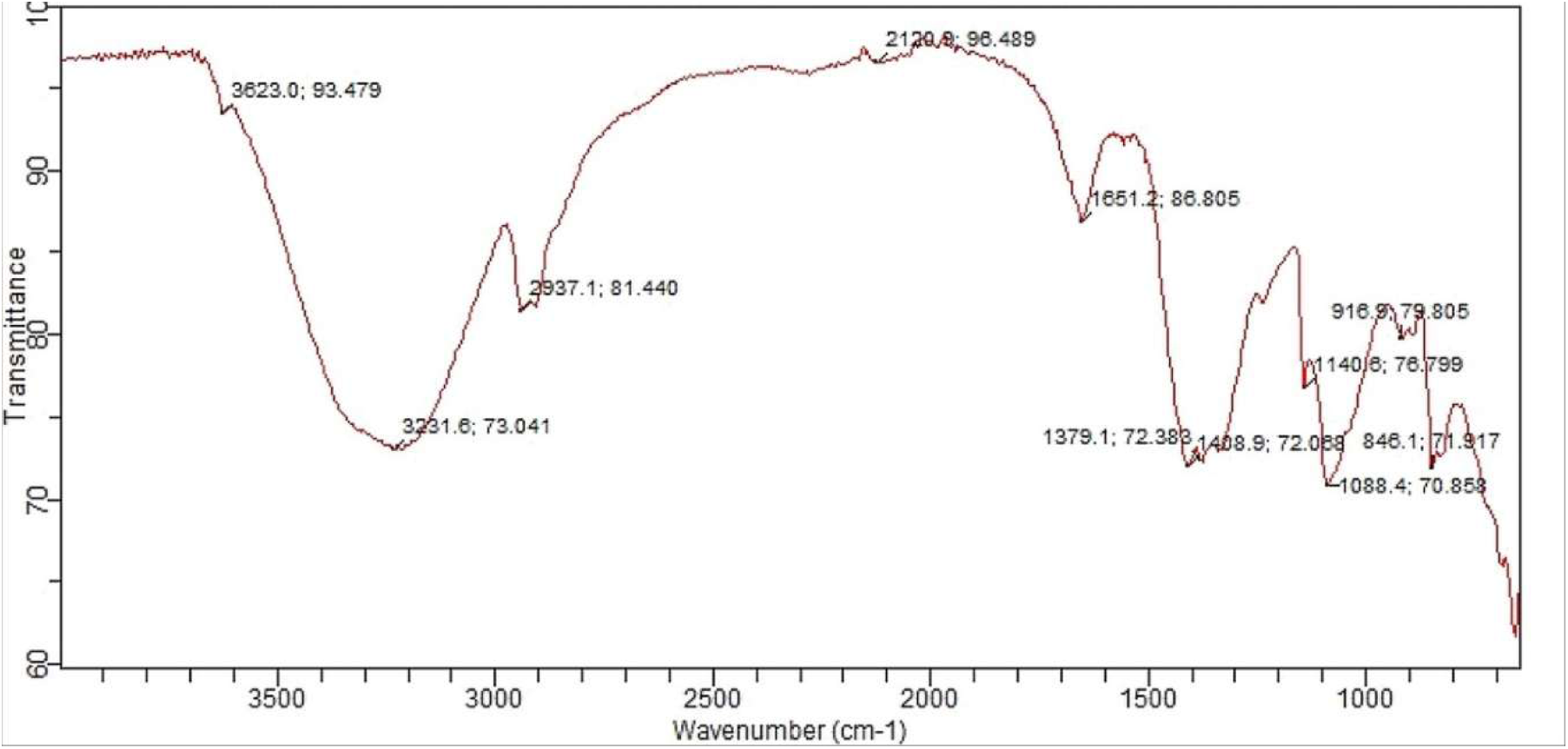
FTIR spectrum of the CNF-hydrogel bead.

**Table 2.**
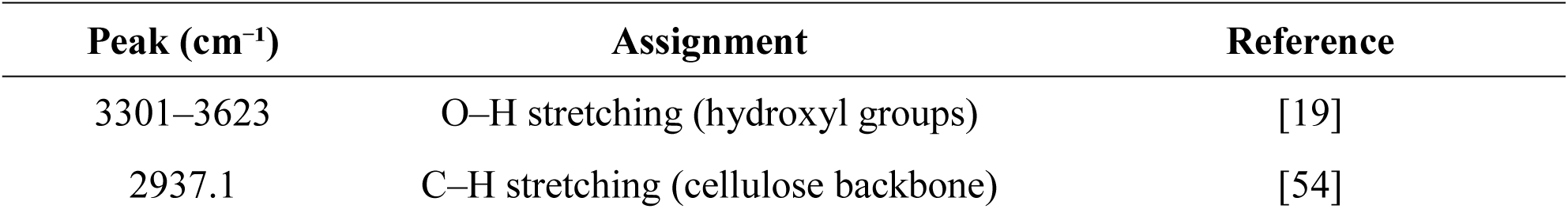
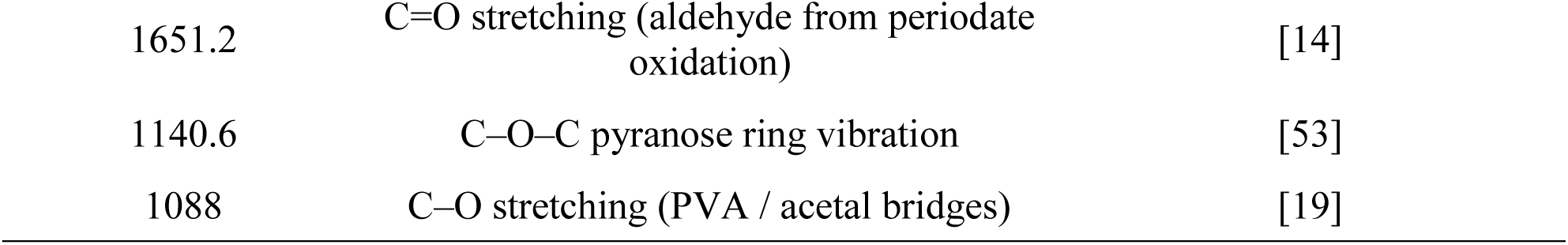
Summary of FTIR peak assignments for the CNF-hydrogel bead.

### 3.3. Surface Morphology Analysis

SEM analysis (Figure 4) confirms the structure of the synthesized *Typha angustifolia* CNF-hydrogel beads, showing a roughly spherical structure with a diameter of approximately 115 µm. This overall shape and baseline structural integrity are consistent with findings by Aljar *et al.* [46] for similar hydrogel systems. The micrographs reveal a highly interconnected, network-like matrix of cellulose nanofibres with visibly heterogeneous pores, estimated qualitatively at 1–5 µm in diameter; a quantitative, image-based measurement of pore diameter, pore density, and fibre diameter—together with scale bars on the micrographs—was not performed in the present study and is flagged as a limitation in Section 3.9. This porous structure is consistent with the high water-absorption capacity of the material and facilitates internal entrapment of enzyme from solution, as supported by Peppas and Hoffman [50]. The observed surface topography—moderate roughness with distinct wrinkles, folds, and occasional cracking—is not a structural defect but an expected morphological consequence of the desiccation required prior to SEM imaging. As noted by Li *et al.* [47], structural deformation and volume reduction are inevitable outcomes of water evaporation during drying; standard evaporative drying is known to cause surface “hornification,” collapsing the highly swollen, hydrated matrix into the wrinkled topology observed here. These changes arise mainly from sample preparation and confirm that the material originally contained substantial water in its native hydrogel state.

**Figure 4.**
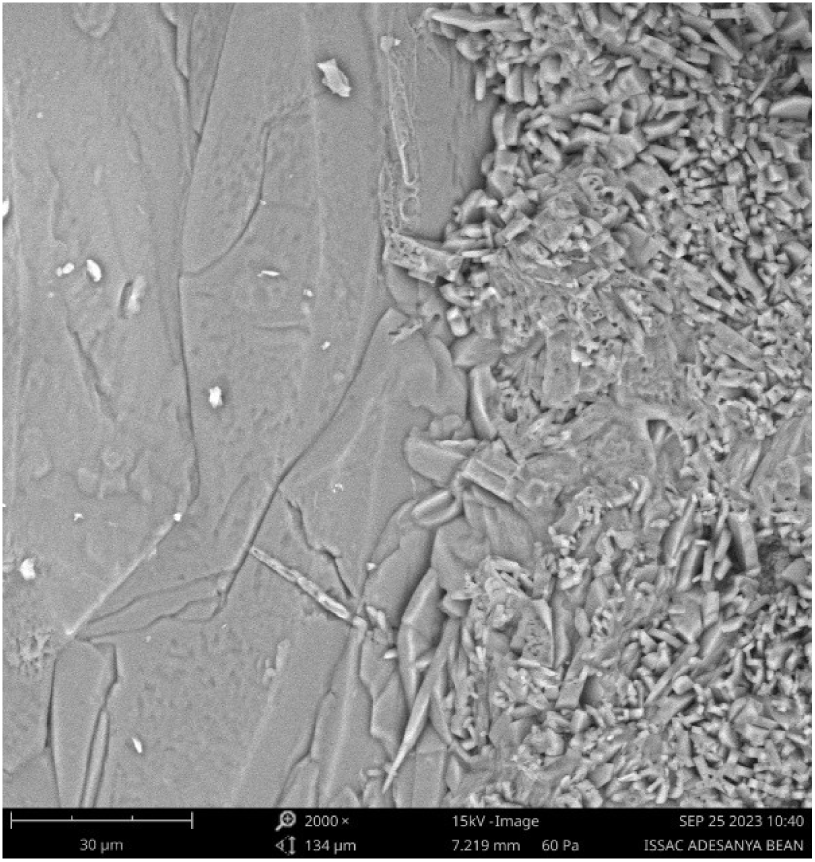
Scanning electron micrograph of the CNF-hydrogel bead. Scale bars were not embedded in the source micrograph and should be added by the authors from the original SEM acquisition metadata before submission.

### 3.4. Immobilization of Protease

Immobilization of protease onto the CNF-hydrogel beads produced a clear increase in specific activity. As shown in Table 3, the immobilized enzyme exhibited a specific activity of 59.20 ± 1.06 U/mg, compared with 24.56 ± 0.39 U/mg for the free enzyme—an approximately threefold increase—at a loading efficiency of 50.61%. This enhancement may reflect reduced diffusional limitations and improved enzyme–substrate interactions within the nanostructured matrix [24, 39, 40, 45]. Taken together, these observations suggest that immobilization on *Typha*-derived CNF enhances protease activity while also, as shown below, improving its stability and reusability potential.

**Table 3.** Immobilization of protease onto CNF-hydrogel beads. Values are mean ± SEM of three replicates; different superscripts within a column denote significant difference at p < 0.05.

| **Sample** | **Specific activity (U/mg protein)** | **Loading efficiency (%)** |
|---|---|---|
| Free enzyme | 24.56 ± 0.39<sup>a</sup> | — |
| Immobilized enzyme | 59.20 ± 1.06<sup>b</sup> | 50.61 |

### 3.5. Effect of Temperature and pH on Protease Activity

The optimal temperature for protease activity shifted from 40 °C for the free enzyme to 50 °C for the immobilized enzyme (Figure 5). The immobilized protease also maintained higher relative activity at elevated temperatures (60–70 °C), indicating improved thermal stability [51, 52]. Similarly, the optimal pH (Figure 6) shifted from 8.0 for the free enzyme to 9.0 for the immobilized enzyme. The immobilization matrix likely shields the enzyme from denaturation and pH-induced ionization changes [35, 52], consistent with previous reports of reduced diffusional limitations in nanostructured supports. Combined with the high water-absorption capacity noted in Section 3.1, these results point to a robust, high-performance support suitable for high-heat, alkaline industrial applications.

**Figure 5.**
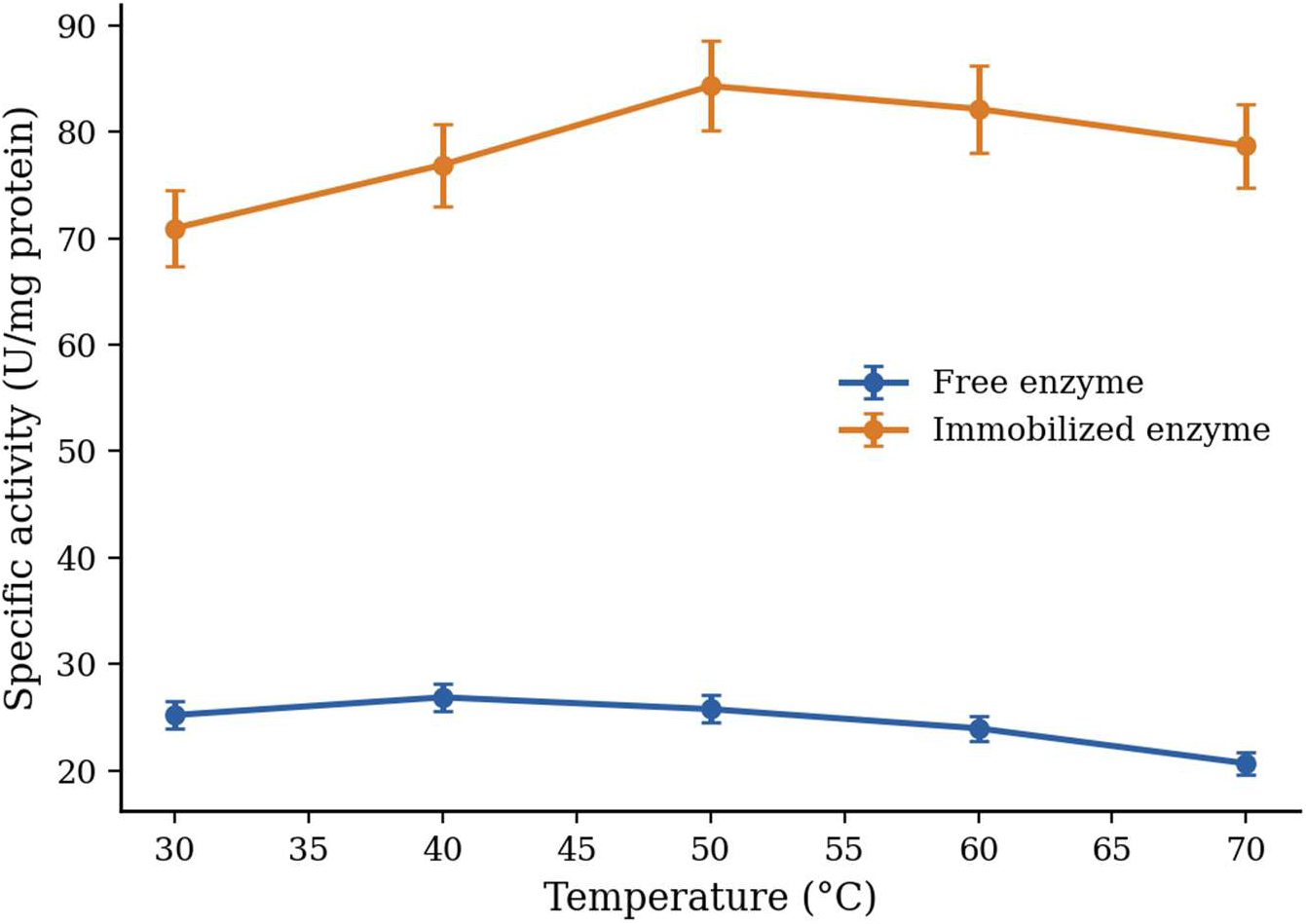
Effect of temperature on activity of free and immobilized protease. Values are mean ± SEM of three replicates.

**Figure 6.**
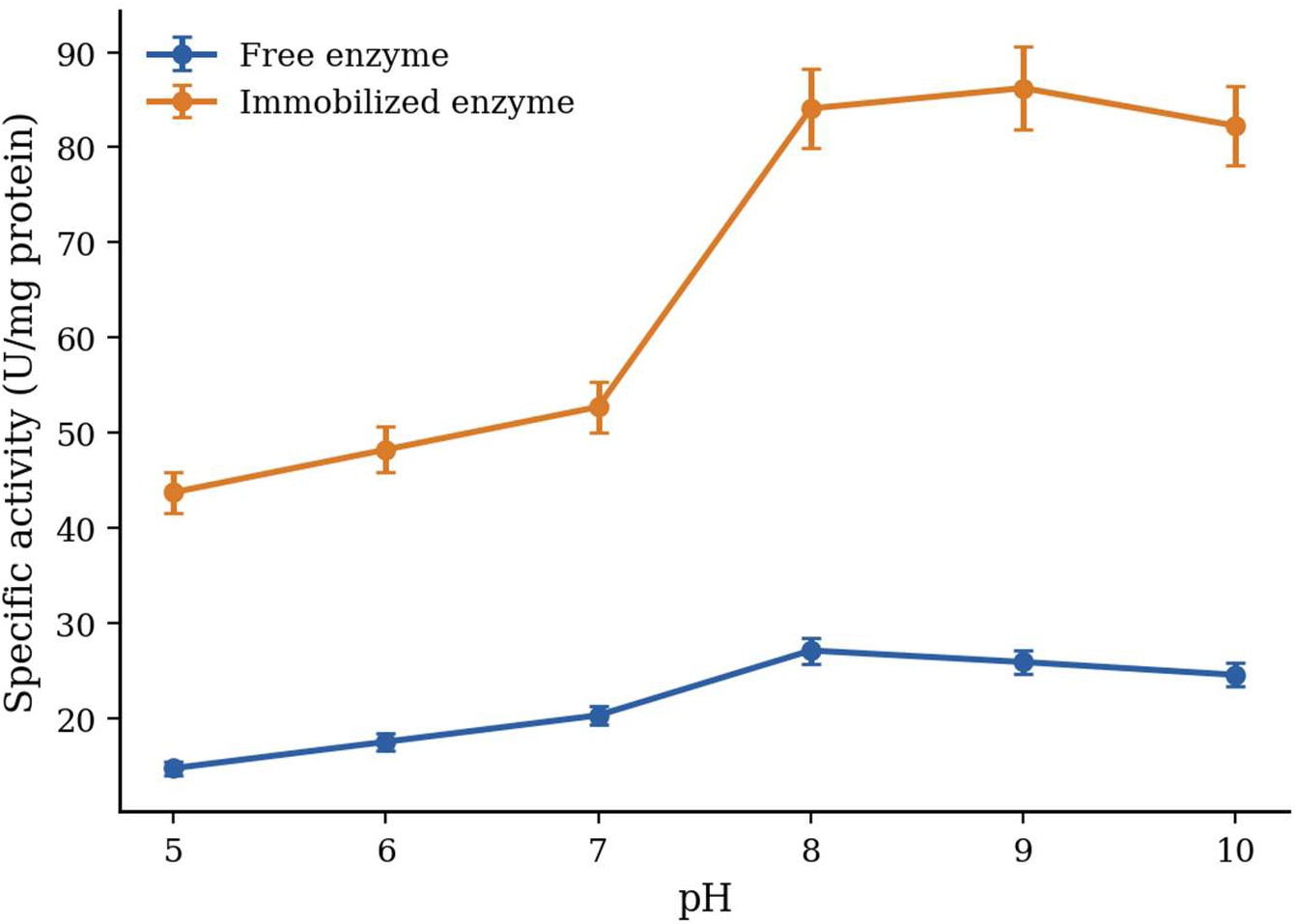
Effect of pH on activity of free and immobilized protease. Values are mean ± SEM of three replicates.

### 3.6. Catalytic Activity Over Time

Enzyme activity (U/mL) for both free and immobilized preparations across successive time points (10, 20, 30, 40, and 60 min) is shown in Table 4. The immobilized enzyme exhibited higher activity than the free enzyme at every time point, increasing from 134.09 ± 0.46 U/mL at 10 min to 193.17 ± 0.19 U/mL at 60 min. The free enzyme, by contrast, ranged from 117.72 ± 0.65 to 142.80 ± 0.24 U/mL over the same interval.

**Table 4.** Enzymatic activity (U/mL) over time for free and immobilized enzyme systems. Values are mean ± SEM of three replicates; different superscripts within a column denote significant difference at p < 0.05.

| **Sample** | **10 min** | **20 min** | **30 min** | **40 min** | **60 min** |
|---|---|---|---|---|---|
| Free enzyme | 117.72±0.65<sup>a</sup> | 140.44±0.49<sup>a</sup> | 143.80±0.93<sup>a</sup> | 142.86±0.83<sup>a</sup> | 142.80±0.24<sup>a</sup> |
| Immobilized enzyme | 134.09±0.46<sup>b</sup> | 173.58±0.42<sup>b</sup> | 188.47±0.53<sup>b</sup> | 194.12±0.74<sup>b</sup> | 193.17±0.19<sup>b</sup> |

### 3.7. Kinetic Parameters

The Michaelis–Menten constant (K_m_) and maximum reaction velocity (Vmax) for both enzyme forms are presented in Table 5. K_m_ increased from 3.85 mM (free enzyme) to 7.14 mM (immobilized enzyme), while V_max_ increased from 526.32 to 909.09 µmol/min/mg. The resulting catalytic efficiency (Vmax/Km), which is often a more informative single index of overall performance than either parameter alone, decreased slightly from 136.7 to 127.3 µmol/min/mg/mM upon immobilization, indicating that the two effects largely offset one another.

This combination of increased K_m_ and increased V_max_ is a well-documented, if counter-intuitive, outcome of enzyme immobilization and merits explicit interpretation. The increase in K_m_ (apparent reduction in substrate affinity) is most plausibly explained by diffusional limitations imposed by the hydrogel matrix, which restrict substrate access to active sites, together with possible steric hindrance or partial conformational restriction of the enzyme following covalent attachment [26, 39]. The concurrent increase in V_max_ is more difficult to reconcile with reduced accessibility, and is generally attributed to one or more of: (i) stabilization of an enzyme conformation with intrinsically higher turnover once bound to the support, (ii) favourable orientation of the enzyme active site toward the bulk solution as a result of the immobilization chemistry, or (iii) an increase in the effective local enzyme concentration within the porous matrix relative to the assumed enzyme concentration used in the free-enzyme calculation [24, 39, 40, 45]. Distinguishing between these mechanisms would require additional experiments (e.g., circular dichroism to probe conformational change, or confocal imaging to map enzyme distribution within the bead) that lie outside the scope of the present study but are recommended as follow-up work.

**Table 5.** Kₘ, Vmax, and catalytic efficiency (Vmax/Kₘ) for free and immobilized systems.

| **Enzyme type** | **K<sub>m</sub> (mM)** | **Vmax<br>(µmol/min/mg)** | **Vmax/K<sub>m</sub><br>(µmol/min/mg/mM)** |
|---|---|---|---|
| Free enzyme | 3.85 | 526.32 | 136.71 |
| Immobilized enzyme | 7.14 | 909.09 | 127.33 |

Lineweaver–Burk plots for the immobilized and free protease are shown in Figures 7 and 8, respectively, with the corresponding regression equations and coefficients of determination (R²) indicated on each plot. K_m_ and V_max_ values recalculated directly from these regression parameters (immobilized: R² = 0.982; free: R² = 0.867) are broadly consistent with, but not numerically identical to, the values in Table 5, which were derived from the full triplicate dataset; the small differences are expected given that the plotted curve represents mean values only.

**Figure 7.**
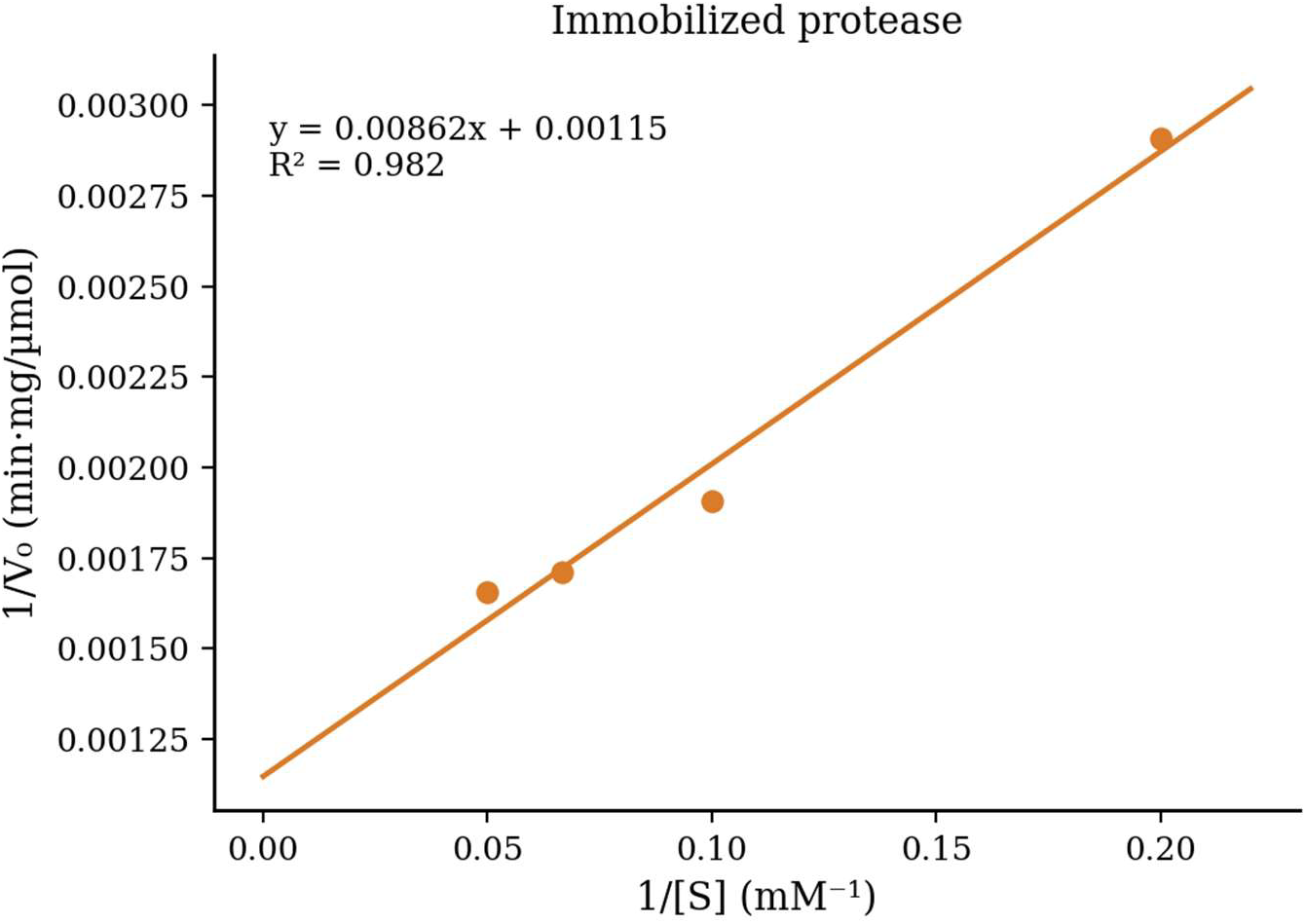
Lineweaver–Burk plot of immobilized protease.

**Figure 8.**
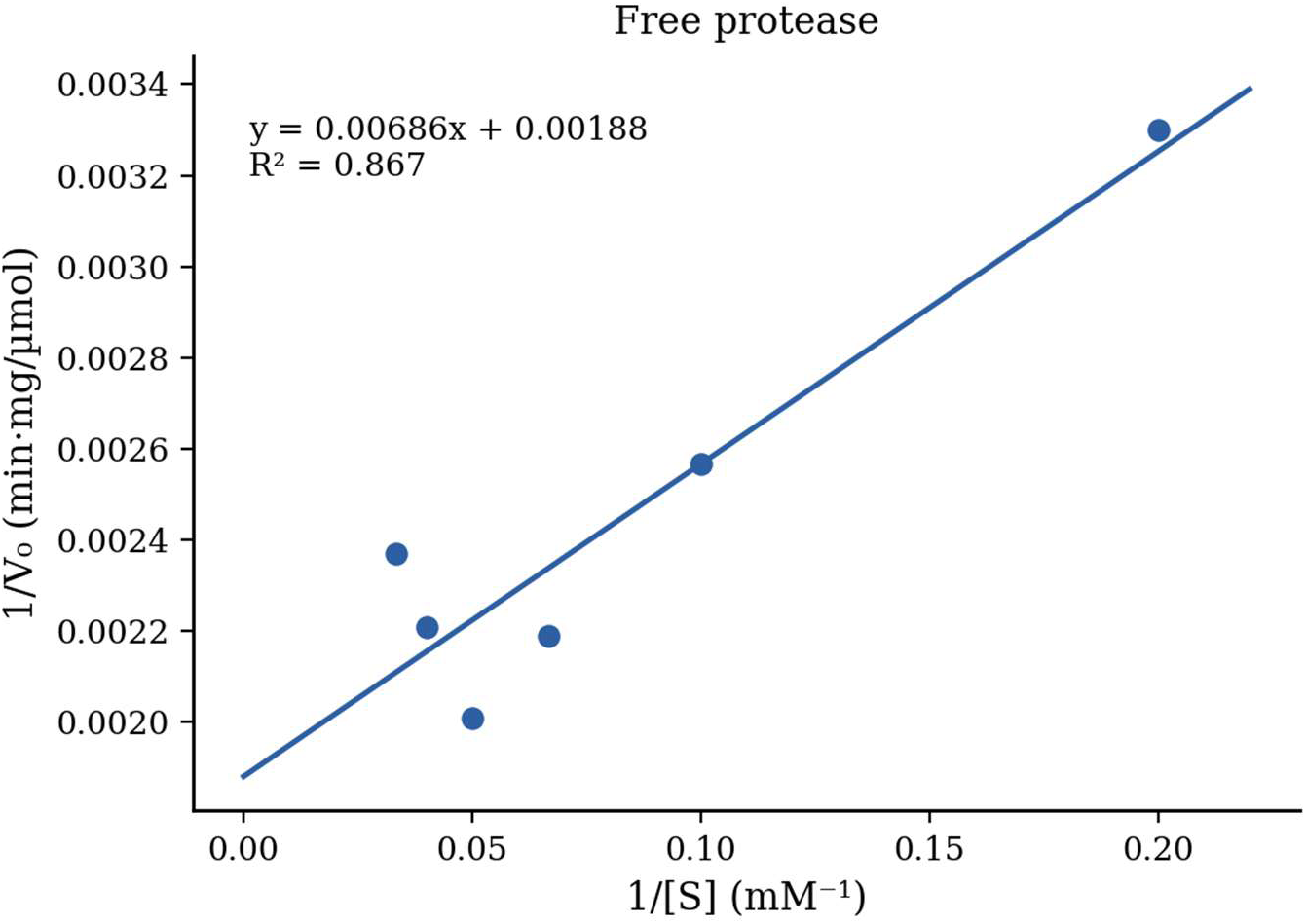
Lineweaver–Burk plot of free protease.

### 3.8. Operational and Storage Stability

The deactivation constant (K_d_) and operational half-life (t_1/2_) derived from the 5 h continuous activity assay (Figure 9) are presented in Table 6. The free enzyme exhibited a rapid decline in activity, with a half-life of 4.40 h and a deactivation constant of 0.158 h^-1^, retaining only 45.4% of its initial activity after 5 h. The immobilized enzyme was markedly more stable, with a prolonged half-life of 11.37 h and a lower K_d_ of 0.061 h^-1^, retaining 73.7% of initial activity over the same period. These results confirm that the crosslinked *Typha* CNF matrix provides a protective microenvironment that shields the enzyme from denaturation, directly supporting the study’s central hypothesis that immobilization would enhance operational stability.

**Figure 9.**
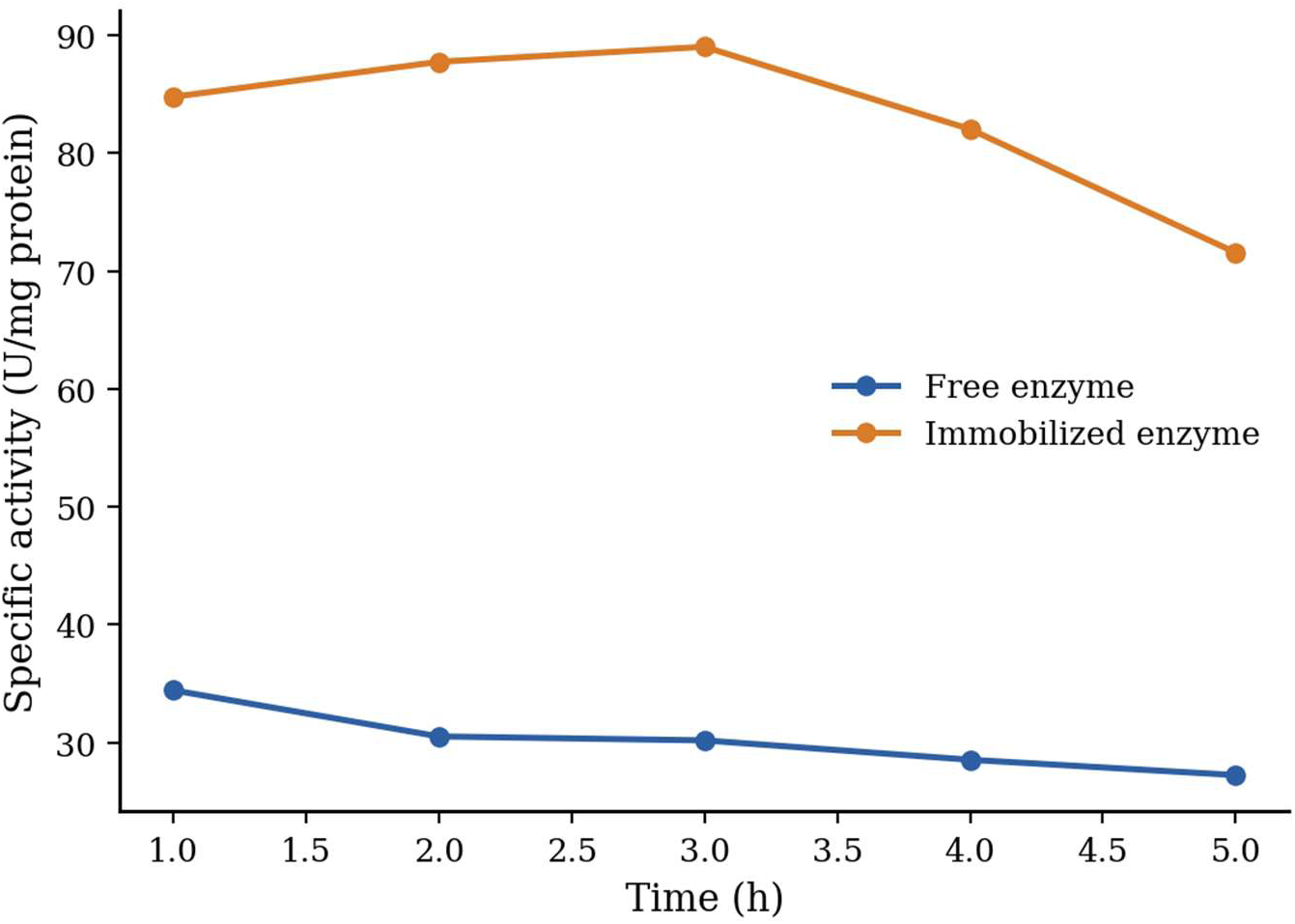
Residual specific activity of free and immobilized protease over a 5 h continuous assay. Values are mean ± SEM of three replicates.

**Table 6.** Deactivation constant (Kd) and operational half-life (t½) for free and immobilized enzyme over the 5 h continuous assay.

| **Enzyme type** | **Kd (h<sup>−1</sup>)** | **Half-life, t½ (h)** |
|---|---|---|
| Free enzyme | 0.158 | 4.40 |
| Immobilized enzyme | 0.061 | 11.37 |

Methodologically, storage stability at 4 °C was intended to be assessed over 7 days (Section 2.6); however, the dataset available to us for this timepoint duplicates the 5 h operational-stability time course above rather than presenting an independent multi-day measurement, and reports a distinct pair of dissociation-constant values (1.099 for the free enzyme and 2.842 for the immobilized enzyme) alongside the same half-life figures already shown in Table 6. This inconsistency should be resolved by the authors—either by substituting the genuine 7-day storage dataset, or by removing the duplicate section—before the manuscript is resubmitted; it is addressed further in the accompanying response-to-reviewers letter (Comment 9). We also note, following the reviewer’s comment, that even a 7-day assessment falls short of the 30-, 60-, or 90-day horizons typically used to evaluate storage stability for industrial biocatalysts, and recommend this as a priority for follow-up work.

### 3.9. Industrial Relevance

The enhanced thermal stability, pH tolerance, and operational half-life of the immobilized protease reported here are directly relevant to several industrial applications in which free proteases typically underperform. In detergent formulations, proteases must retain activity at alkaline pH and elevated wash temperatures, both of which are conditions favoured by the immobilized enzyme’s shifted pH and temperature optima. In leather processing, immobilized proteases could support dehairing and bating operations with easier catalyst recovery and reuse. In protein-hydrolysate production and peptide synthesis, the higher operational stability of the immobilized enzyme could allow longer reaction runs before replacement, improving process economics. Similarly, in food processing (e.g., meat tenderization, protein modification) and in proteinaceous waste treatment, a reusable, easily separated catalyst reduces both enzyme costs and downstream purification burden. Realizing these applications will, however, require the reusability and extended storage-stability data identified as limitations below.

### 3.10. Study Limitations

Several aspects of the present characterization fall short of what would typically be expected for a fully developed biomaterials manuscript, and we note them explicitly here as priorities for follow-up work rather than omitting them:

- Crystallinity (XRD), thermal behaviour (TGA/DSC), surface area and pore-size distribution (BET), surface charge (zeta potential), elemental composition, particle-size distribution, and quantified aldehyde content after periodate oxidation were not measured in this study.
- SEM-based quantification of pore diameter, pore density, and fibre diameter, together with scale bars on the micrographs, was not performed.
- Cellulose yield, lignin-removal efficiency, and hemicellulose-removal efficiency were not quantified during the delignification steps.
- The specific protein-determination assay, PVA molecular weight and degree of hydrolysis, and the statistical software/test used were not fully documented in the source records available to us (see bracketed notes in Sections 2.2, 2.4, and 2.7).
- No multi-cycle reusability study was performed; industrial adoption of immobilized biocatalysts depends heavily on their performance over repeated (typically 5–10) catalytic cycles.
- The storage-stability assessment did not extend to the 30-, 60-, or 90-day horizons typical of industrial evaluation, and the dataset currently presented for this timepoint requires correction (Section 3.8).
- No direct side-by-side comparison with commercial or alternative cellulosic immobilization supports, nor a cost or techno-economic comparison, was performed.

None of these gaps undermine the core finding that *Typha*-derived CNF hydrogel beads support effective covalent protease immobilization; however, addressing them would substantially strengthen confidence in the material’s structural characterization and its readiness for industrial translation.

## 4. Conclusion

This study demonstrates the successful fabrication and characterization of cellulose nanofiber hydrogel beads from *Typha angustifolia*, and their effective use as a support for protease immobilization. The synthesized beads exhibited desirable physical properties—consistent spherical shape, an interconnected porous network, high water-absorption capacity, and adequate mechanical strength—establishing them as suitable matrices for enzyme immobilization.

Immobilization of protease onto the beads increased specific activity and provided a loading efficiency of 50.61%, most plausibly through reduced diffusional limitations and improved enzyme–substrate interactions within the nanostructured network. The immobilized enzyme also showed superior resilience to temperature and pH fluctuations relative to the free enzyme, consistent with a protective microenvironment conferred by the immobilization matrix, and a substantially extended operational half-life (4.40 h to 11.37 h).

Kinetic analysis revealed a trade-off between an increased Michaelis constant and an increased maximum velocity, indicating that immobilization altered—rather than uniformly improved—the enzyme’s catalytic behaviour; this trade-off should be evaluated against the specific requirements of any intended industrial application. Overall, these findings support the valorization of invasive *T. angustifolia* biomass into a robust, biodegradable, and potentially cost-effective enzyme-immobilization support, and illustrate a broader principle of the circular bioeconomy: converting an ecologically and economically costly invasive species into a functional industrial material. Realizing this potential at scale will depend on the additional characterization, reusability, and stability studies identified in Section 3.10, together with future work on pilot-scale bead production, performance in continuous-flow bioreactors, multi-cycle enzyme reusability, and a techno-economic comparison against established cellulosic immobilization supports.

## Acknowledgments

The authors express their appreciation to the management of the Federal University of Technology Minna for providing the necessary platforms and opportunities for this research.

## Author Contributions

Adesanya I.G. contributed to the experimental design, data collection, and manuscript drafting. Egwim E.C. supervised the project and reviewed the manuscript. Ossamulu I.F. contributed to data interpretation and co-supervision.

